# Retrospective analysis of enhancer activity and transcriptome history

**DOI:** 10.1101/2021.09.07.459233

**Authors:** Ruben Boers, Joachim Boers, Beatrice Tan, Evelyne Wassenaar, Erlantz Gonzalez Sanchez, Esther Sleddens, Yasha Tenhagen, Marieke E. van Leeuwen, Eskeatnaf Mulugeta, Joop Laven, Menno Creyghton, Willy Baarends, Wilfred F. J. van IJcken, Joost Gribnau

## Abstract

Cell state changes in development and disease are controlled by gene regulatory networks, the dynamics of which are difficult to track in real time. Here, we utilize an inducible DCM-RNA-polymerase-subunit-b fusion protein, to label active genes and enhancers with a bacterial methylation mark that does not affect gene transcription and is propagated in S-phase. We applied this DCM-time-machine (DCM-TM) technology to study intestinal homeostasis, following enterocyte differentiation back in time, revealing rapid and simultaneous activation of enhancers and nearby genes during intestinal stem cell (ISC) differentiation. We provide new insights in the absorptive-secretory lineage decision in ISC differentiation, and show that ISCs retain a unique chromatin landscape required to maintain ISC identity and delineate future expression of differentiation associated genes. DCM-TM has wide applicability in tracking cell states, providing new insights in the regulatory networks underlying cell state changes in development and differentiation.

## Introduction

Embryonic development and cell differentiation are intricate processes directed by cross talk between cells that affect cell fate decisions and the establishment of cell type specific gene expression programs (Bradner et al., 2017; Lee and Young, 2013; Stadhouders et al., 2019). Lineage tracing studies have been crucial to understand these processes. Initial studies applied light microscopy to follow cleavage divisions, but since then bar-coding, cre-lox and other genetic systems have been utilized to mark precursors or progenitors for readout at later stages of development or differentiation (Alemany et al., 2018). The present advance of the single cell RNA sequencing (scRNA-seq) technology provides a wealth of expression data that can be used to predict developmental trajectories *in silico* and can be linked to genetic lineage tracing techniques to rebuild lineage trees (Bowling et al., 2020; Herman et al., 2018; Schiebinger et al., 2019).

Application of these tracing technologies to study the epithelium of the small intestine provided critical insights in homeostasis and regeneration. Turnover of this epithelium happens within 7 days and starts with division of the intestinal stem cell (ISC) located at the bottom of the intestinal crypt (Beumer and Clevers, 2020). ISCs give rise to progenitors that divide moving up the intestinal crypt meanwhile committing to absorptive or secretory lineage. Absorptive progenitors mature into enterocytes, whereas secretory progenitors give rise to Paneth, tuft, entero-endocrine and goblet cells. ISCs are flanked by Paneth cells that provide Wnt, Notch and EGF signals required for self-renewal. Loss of ISC-Paneth cell contact facilitates cell differentiation, aided by BMP signalling that further supports maturation of differentiated cell types. Notch signalling also plays a crucial role in lineage commitment remaining high in absorptive progenitors and is downregulated in secretory progenitors. Lineage tracing and scRNA-seq experiments have been instrumental in identification and characterisation of the crypt based columnar cell as the ISC (Barker et al., 2007), but also showed that several other cell types including entero-endocrine, Paneth and immature enterocytes provide a reservoir of cells that can replenish the ISC niche in injury induced regeneration (Tetteh et al., 2016; Yan et al., 2017; Yu et al., 2018).

While these examples highlight the successful application of lineage tracing and scRNA-seq technologies to build relationships between cellular trajectories they cannot keep track of cell state changes following this trajectory and provide limited depth and temporal information with respect to gene expression changes (Baron and van Oudenaarden, 2019). To facilitate whole genome cell state tracing, we developed a system to epigenetically tag transcribed genes to be examined at later stages of development or differentiation. We made use of a fusion between DCM and RNA-polymerase-2-subunit-b to epigenetically label gene bodies of transcribed genes. DCM methylation of C_me_C(A/T)GG penta-nucleotides is a bacterial form of cytosine methylation only detected at very low levels in most mammalian cell types, but is maintained when introduced on transgenes in somatic cells without affecting transgene expression (Clark et al., 1995). Our study demonstrates that DCM-TM marks both active genes as well as enhancers, both of which are occupied by RNA-polymerase-2 and confirms that DCM methylation is propagated to daughter cells with limited effect on gene expression. Thus DCM-TM provides a powerful technology to trace genome wide gene transcription and enhancer activity back in time without relying on in silico assumptions. We applied DCM-TM to study homeostasis in the small intestine, generating gene and enhancer activity maps that trace the ISC state to the enterocyte state. We discovered that gene and enhancer activity changes during enterocyte differentiation are not mediated by heterochromatin changes and show that the H2A variant H2A.Z is preloaded at ISC enhancers that will become activated in the enterocyte. Application of DCM-TM also indicated that commitment of progenitors to the absorptive lineage is a one-way event which does not involve a temporarily dynamic absorptive-secretory intermediate state.

## Results

### DCM-Polr2b marks active genes and enhancers

To develop a gene activity tagging system we fused DCM to the N-terminal end of mouse RNA-polymerase-2 subunit-b (*Polr2b*, Fig 1a), and introduced this DCM-Polr2b fusion gene into the *Col1a1* locus, in an ESC line harbouring the m2rtTA trans-activator expressed from the *Rosa26* locus (Sup Fig 1a-b)(Beard et al., 2006). Addition of doxycycline leads to expression of the fusion protein at levels comparable to endogenous POLR2B, and expression is depleted 24 hours after removal of doxycycline (Sup Fig 1c-d). To detect DCM methylation we developed methylated DNA sequencing (MeD-seq), a technology based on LpnPI mediated digestion of CpG and DCM methylated target sites resulting in 32 base pair fragments that are sequenced (Fig 1a)(Boers et al., 2018). LpnPI recognizes 50% of all methylated CpG di-nucleotides (C_me_CG, _me_CGG, and G_me_CGC), as well as DCM methylated C_me_C(A/T)GG penta-nucleotides.

**Figure 1.**
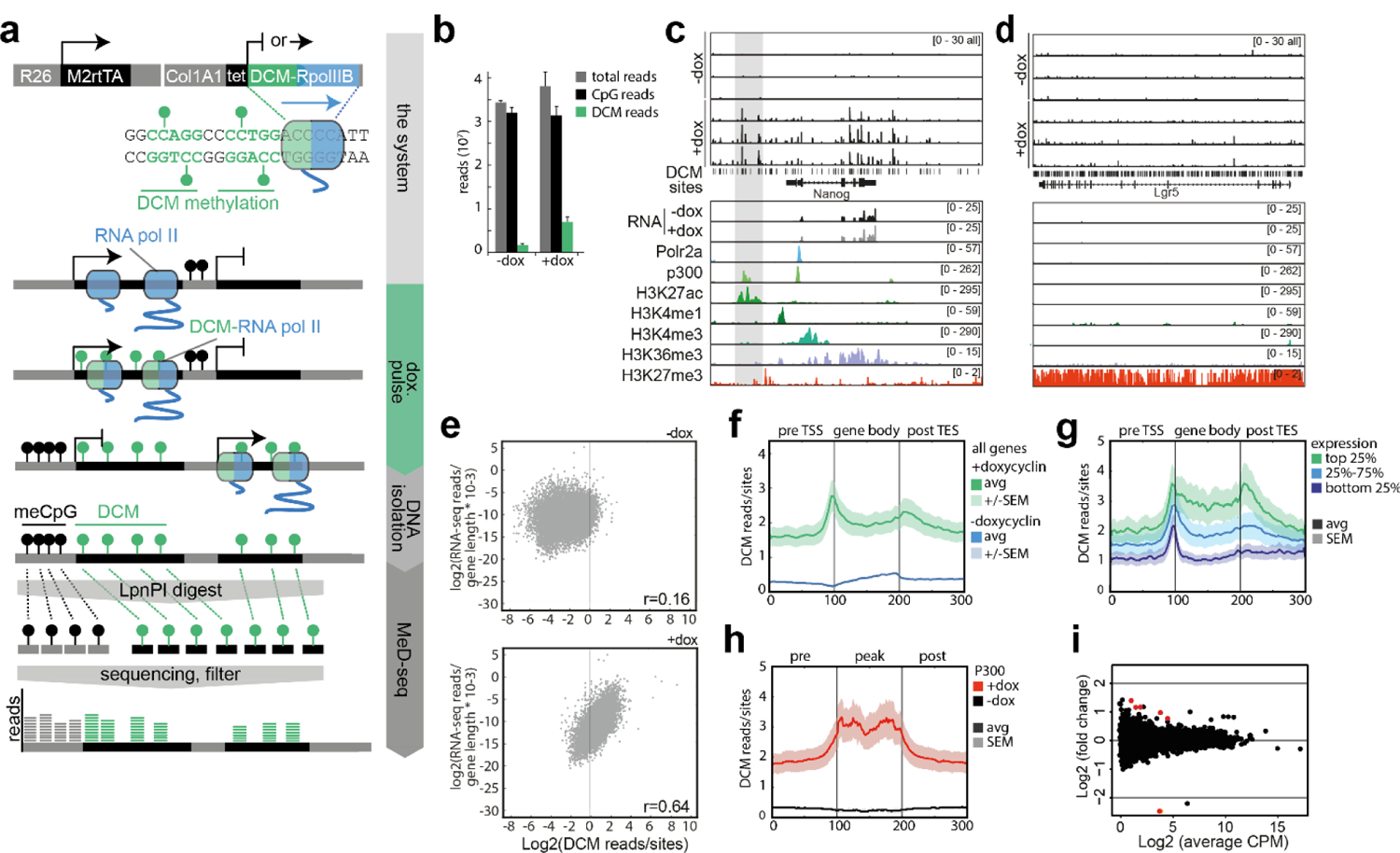
The DCM-time-machine in embryonic stem cells. (a) Overview of the DCM-TM and MeD-seq pipelines. (b) Induction of DCM labelling measured 5 days after start of dox treatment. (c,d) Genome browser view of DCM specific MeD-seq reads (n=3), RNA-seq (+/- dox) and ChIP-seq tracks (ENCODE) in the *Nanog* (c, enhancer indicated in grey) and *Lgr5* (d) loci. (e) Scatter plot displaying RNA-seq gene expression level in relation to DCM read count per gene before (top panel) and after dox induction (bottom panel). (f) Gene meta-analysis showing binned distribution of DCM reads overlapping the gene body before (blue) and after (green) 5 days of dox treatment. The average (avg) and standard error of the mean (SEM) are depicted with a darker line and lighter region, respectively. (g) Gene meta-analysis showing distribution of DCM reads of expressed genes split in three clusters based on expression (top 25%, 25%-75% and bottom 25%) after 5 days of dox treatment. (h) DCM read meta-analysis showing binned distribution of DCM read counts over p300 positive genomic regions and 1kb proximal and distal flanking regions. (i) RNA-seq analysis comparing average gene expression values before and 5 days after dox induction (genes indicated in red show significant expression change).

Addition of doxycyclin (dox) to DCM-Polr2b:m2rtTA ESCs for five days resulted in a 5-fold induction of DCM methylation genome wide (Fig 1b, Sup Fig 1e, Sup Table 1). DCM methylation clearly increased in gene bodies of genes expressed in ESCs (*Nanog*, *Zfp42*, *ActB*), while no accumulation was observed in genes not expressed in ESCs (*Lgr5*, *Alpi*, Fig 1c-d, Sup Fig 1f). The distribution of DCM sites is clearly different from CpG sites, showing no clear enrichment at the TSS (Sup Fig 1f). In genes with at least 10 DCM sites, uninduced gene body DCM methylation displayed little correlation with gene expression, while after dox induction this correlation became robust (Fig 1e, Sup Fig 1g). Gene-meta-analysis indicated that the DCM methylation profile before dox induction resembled the distribution of CpG methylation that was present in gene bodies of active genes and possibly introduced as an accidental by-product of CpG methylation (Fig 1f, Sup Fig 1k-m) (Arand et al., 2012). After induction, the DCM methylation profile displayed increased DCM methylation at the transcription start sites (TSS), gene body and transcription end sites (TES), with a direct relationship between gene expression and DCM methylation levels (Fig 1f-g).

Comparison with published ChIP-seq data confirmed accumulation of DCM methylation at the TSS and gene body (H3K36me3) after dox induction. In addition, we found DCM methylation to accumulate at regions marked by enhancer specific modifications or protein recruitment (P300, H3K27Ac and H3K4me1, Fig 1c-d, h, and Sup Fig 2a-b) which is consistent with Pol2 recruitment occurring at enhancers (Kim et al., 2010). Whole genome differentially methylated region (DMR) calling (+dox vs – dox, Mann-Whitney significance test) identified 5,973 regions, displaying significantly increased DCM methylation levels, enriched for enhancer specific histone modifications, DNase sensitivity, and pluripotency factor binding (Sup Fig 2c). DCM methylation levels were significantly elevated in genes located in closest proximity to these intergenic DCM DMRs, and enhancer density was proportional to activity of the closest gene (Sup Fig 2d-e)(Kim et al., 2010). Only 6 genes responded, with a change in gene expression, following induction of the DCM-Polr2b fusion gene (Fig 1i), confirming an earlier report that DCM directed methylation of gene bodies has little effect on gene expression (Clark et al., 1995). These results indicate that DCM-Polr2b is functional and labels active genes and enhancers with minimal effect on gene expression.

**Figure 2.**
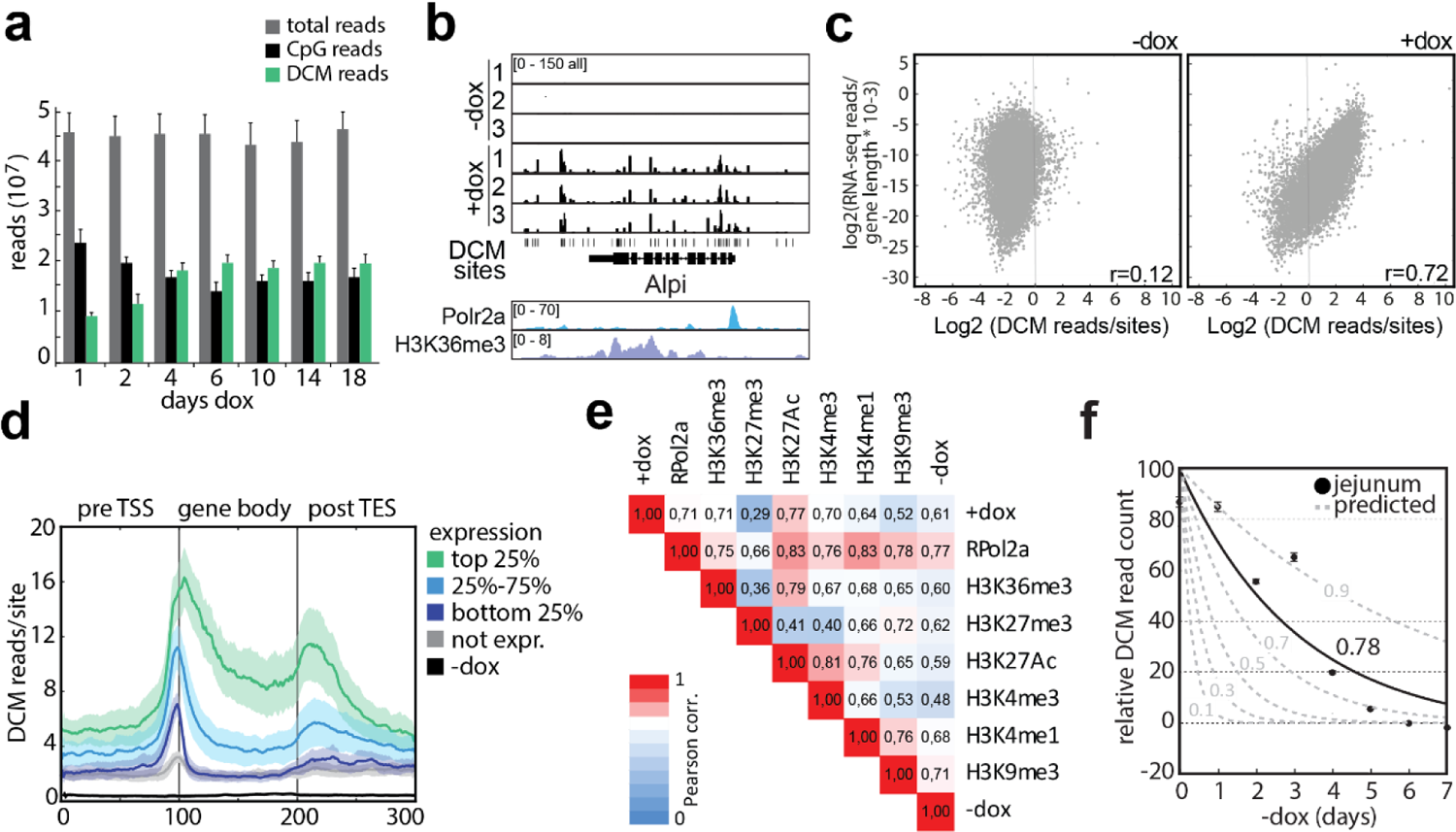
DCM labelling and propagation in the small intestine. (a) Genome wide DCM and CpG methylation at different time points after start of dox induction. (b) Genome browser view of *Alpi* locus showing DCM specific MeD-seq reads before and after 1 day dox treatment. Polr2a and H3K36me3 ChIP-seq tracks from ENCODE are shown below. (c) Scatter plot displaying RNA-seq gene expression level in relation to DCM read count per gene before (left panel) and after 10 days of dox induction (right panel) in epithelium of jejunum. (d) Gene meta-analysis showing distribution of DCM reads in the top 25%, 25%-75% and bottom 25% expressed genes after 3 days of dox treatment. (e) Pearson correlation analysis comparing DCM and ChIP-seq read count distribution. (f) Relative DCM read count measurement at different time points after withdrawal of dox and the estimated propagation rate per cell division in jejunum. The dotted grey lines show the estimated curves based on several different propagation rates, whereas the propagation rate fitted to the DCM data is plotted in black.

### DCM methylation propagation in vivo

To monitor accumulation, maintenance and propagation of DCM methylation *in vivo* we generated DCM-Polr2b transgenic mice. Total epithelium of jejunum was isolated through mechanical shearing from transgenic mice treated with doxycyclin from day 0 through day 18. MeD-seq analysis indicated induced DCM methylation to plateau around day 6 with a >25-fold induction over endogenous DCM methylation levels (Fig 2a-b, Sup Table 1, Sup Fig 3a). Similar as observed in ESCs DCM methylation of TSS, gene body and TES, correlated with gene expression level, Polr2a binding, H3K36me3 deposition (transcribed genes), and H3K27Ac enriched regions (active promoters and enhancers, Fig 2b-e). In addition, DCM gene density distribution was clearly distinct from CpG methylation (Sup Fig 3b). To determine the *in vivo* DCM methylation propagation rate in jejunum, mice were administered dox for 7 days, followed by a chase period of 0-7 days after which DCM/CpG methylation ratios were established in triplicate by MeD-seq. With an estimated average cell cycle of 18.3 hours, and taking into account that MeD-seq analysis does not detect hemi-methylated templates, DCM propagation was estimated to be 0.78 per cell division (Fig 2f)(Parker et al., 2017). We found no difference in the propagation rate for specific genomic regions such as gene bodies, exons, introns, CpG islands, or intergenic regions (Sup Fig 3c). Longer chase periods for up to two months after induction, still revealed DCM methylation gene density profiles that were clearly distinct from controls (Sup Fig 3d) suggesting our method is compatible with cell state tracing across longer temporal windows.

**Figure 3.**
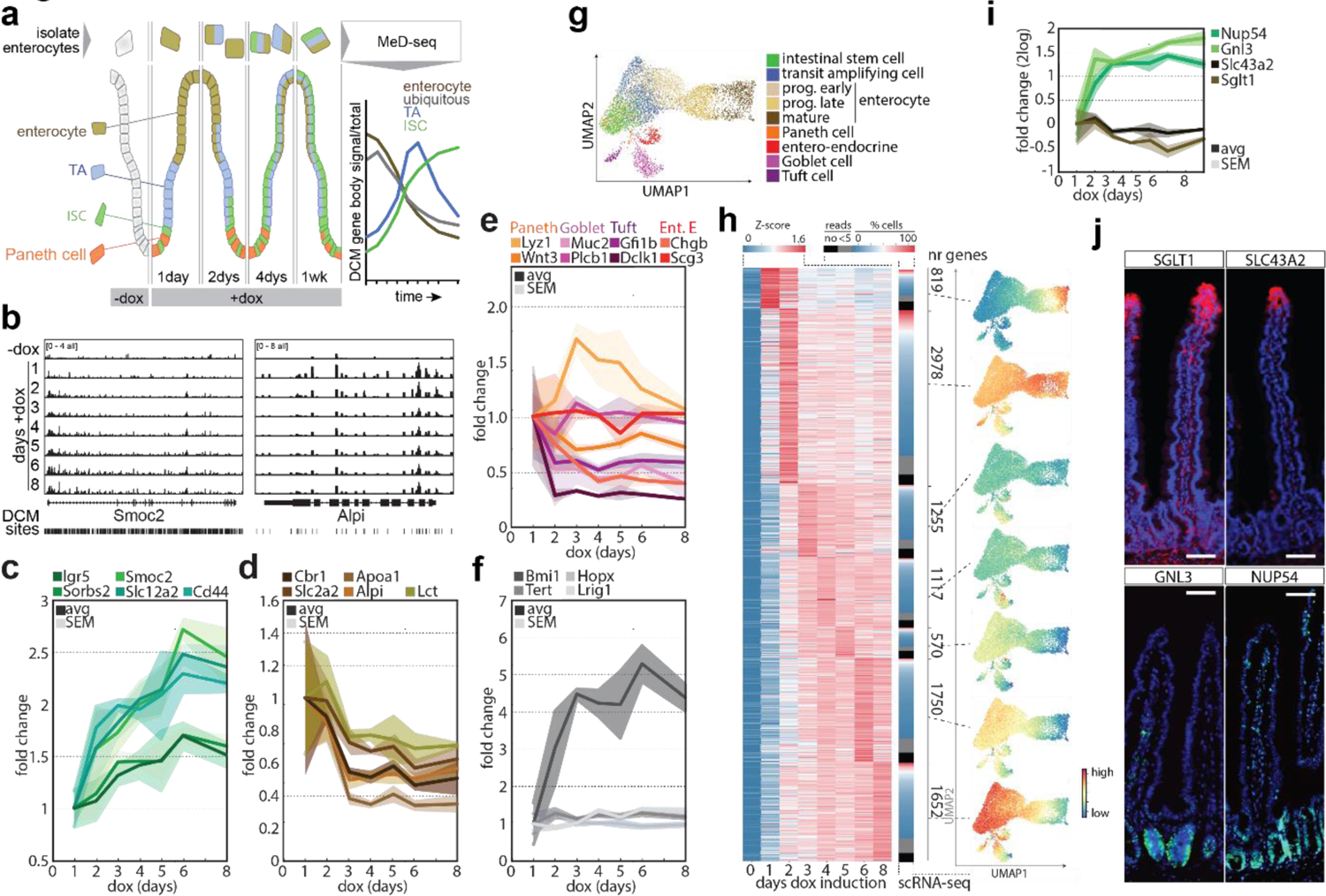
DCM-Rpol2b labelling reveals gene activity maps from ISC to enterocyte. (a) Overview of experimental procedure: mice are labelled with dox and sacrificed at different time points to isolate Epcam+/SLC2A2+ positive enterocytes that are subjected to MeD-seq. ISC, TA, enterocyte and ubiquitously expressed genes are expected to display different dynamic behaviour in time. (b) Genome browser view of the average normalized MeD-seq DCM reads (n=3) in the *Lgr5* and *Alpi* loci at different time points after start of dox treatment. (c-f) DCM labelling (fold change in DCM reads relative to total and normalized to T=1d) of ISC (c), enterocyte (d), Paneth, Goblet, Tuft, Entero-endocrine (e) and +4 cell (f) specific genes. (g) UMAP of jejunum scRNA-seq data showing clusters annotated as specific cell types. (h) DCM labelling of all significantly labelled genes (negative (T=0) samples compared to all days after start of dox treatment) clustered according to the maximum DCM signal, capture of clustered genes by scRNA-seq (for each gene with >5 reads percentage of cells with signal is indicated), and average expression of clustered genes in UMAP shown in (g). (i-j) DCM labelling (i) and validation by immuno-cytochemistry (j) of SGLT1, SLC43A2, GNL3 and NUP54 expression (FITC, DNA is DAPI stained, scale bar: 50μm).

### Gene activity dynamics in ISC to enterocyte differentiation

As turnover of the epithelium is very high in the intestine, we tested our DCM-TM technology following the differentiation of ICSs into enterocytes that are eventually shed from the top of the villi. We isolated enterocytes, through sequential purification of small intestinal epithelium, followed by FACS sorting of Epcam+/SLC2A2+(GLUT2) enterocytes (Fig 3a, Sup Fig 4a). Comparison of RNA-seq data of isolated cells with published scRNA-seq data obtained from intestinal epithelium confirmed proper isolation of proximal enterocytes (Sup Fig 4a-c)(Haber et al., 2017). Subsequently, the DCM-Polr2b fusion gene was induced with doxycycline and enterocytes were isolated at different time points across an eight-day window, followed by MeD-seq and normalization for DCM induction efficiency (Sup Fig 4d, Sup Table 2). In this setting turn-over of the fusion protein is not required and non-dividing cells do not affect the assay. As time passes, DCM methylation levels of stem cell specific genes are expected to increase relative to the total pool of DCM sequencing reads as their profile will be propagated in the transit amplifying (TA) cells and enterocytes (Fig 3a). In contrast, DCM reads from enterocyte specific genes as well as ubiquitous expressed genes are expected to decline over time as they are shed from the top of the villi. Indeed, stem cell markers *Lgr5*, *Sorbs2*, *Smoc2*, and *Slc12a2*, Wnt target gene *Cd44*, and ephrin receptors *EphB2* and *EphB3* displayed an increase in DCM gene body labelling reaching a maximum signal at day 6 (Fig 3b-c, Sup Fig 4e). Ephrin receptor ligands expressed in the villus, *Efnb1* and *Efnb2*, and enterocyte markers *Cbr1*, *Slc2a2*, *Apoa1*, *Alpi* and *Lct* displayed a decrease in DCM methylation (Fig 3b,d, Sup Fig 4e). Ubiquitously expressed genes resembled enterocyte specific genes but with slower kinetics and less dynamic behaviour (Sup Fig 4f). Finally, genes associated with other differentiated cell types or cell types implicated in injury-induced plasticity, including the Paneth, goblet, tuft, entero-endocrine and +4 cell, either did not increase over time, or remained below background levels, indicating no role for these cell types in intestinal homeostasis in the measured time span (Fig 3e-f). One clear exception was *Bmi1*, contrasting with other +4 cell markers in behaviour by resembling other ISC genes. This emphasizes a role for *Bmi1* in ISC homeostasis in line with studies indicating *Bmi1* to be essential for ISC maintenance and intestinal homeostasis (Lopez-Arribillaga et al., 2015).

**Figure 4.**
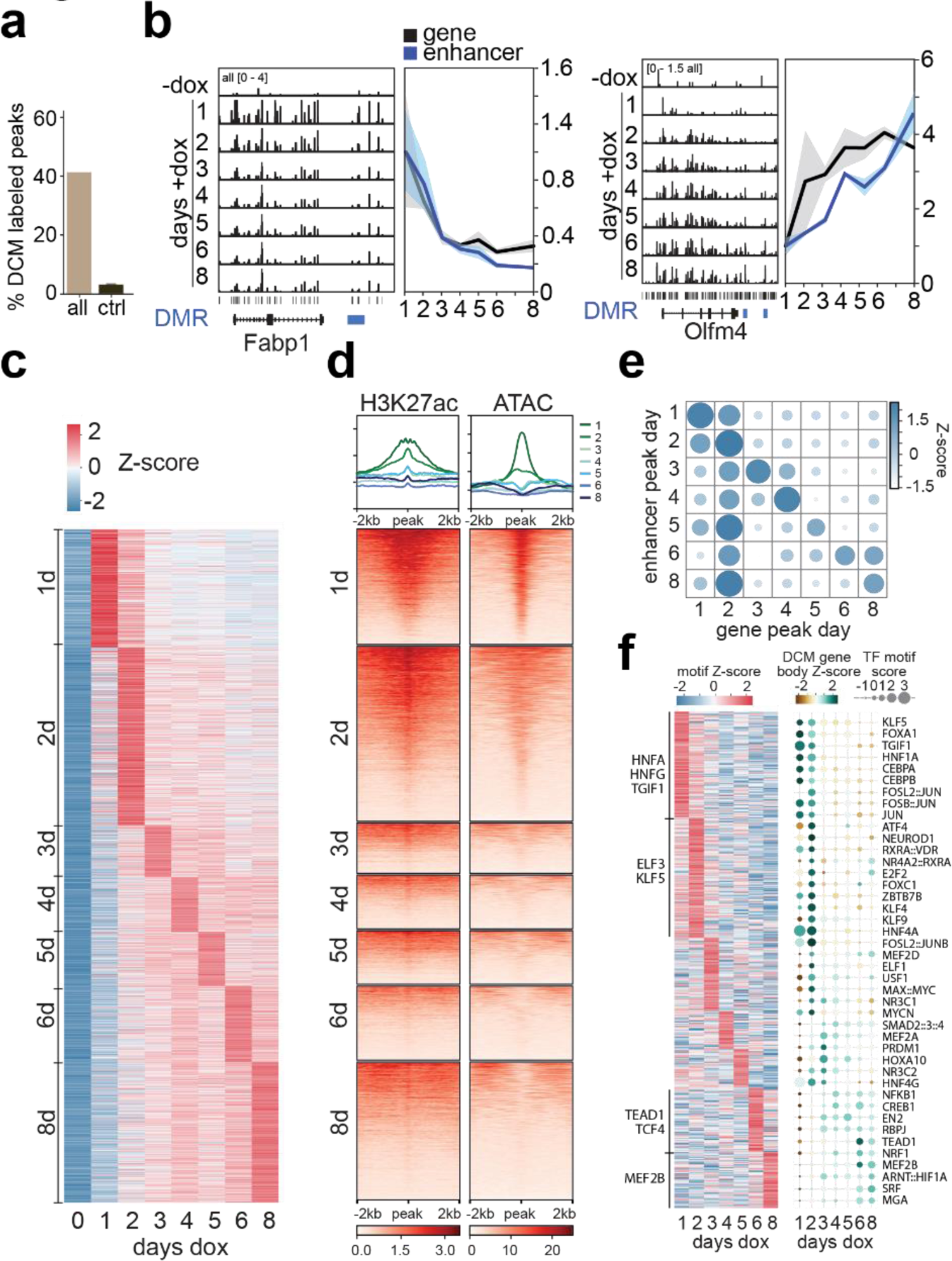
Temporal changes in TF and enhancer activity from ISC to enterocyte. (a) Percentage of H3K27ac peaks that are labelled by DCM (i.e. peaks with ≥ 1 significant DCM site <750 bp from peak). Random control based on 100 sets of reshuffled H3K27ac peaks is added to show expected random overlap (mean ± SD). (b) Left panels display genome browser view showing DCM labelling of enterocyte specific (*Fabp1*) and ISC specific (*Olfm4*) genes with nearby enhancers (marked in blue) showing coordinated behaviour in time. Right panels show the average profiles over time for each gene and the average of the closest significant DCM sites. (c) Heatmap of DCM labelling of enhancers (Z-scores of mean normalized DCM reads). (d) Heatmap showing H3K27ac ChIP-seq and ATAC-seq overlap with the regions around enhancer DMRs peaking at different days of dox induction. Each profile plot has the same y-axis range as its corresponding heatmap. (e) Correlation between gene peak day and peak day of close-by enhancers (z-score of proportion of enhancers per day). (f) Heatmap showing TF motif dynamics observed in intergenic DCM DMRs in time (left) and combined analysis of motif enrichment and DCM gene body labelling dynamics of TFs displaying a positive correlation in time (right).

To generate temporal gene activity maps throughout ISC to enterocyte differentiation, genes with a DCM signal significantly higher than background were clustered according to their temporal signal strength using maximum signal day, and average expression of the different gene clusters was displayed on a UMAP that was based on scRNA-seq data from intestinal epithelium (Fig 3g-h)(Haber et al., 2017). This analysis showed that genes with a temporal methylation profile peaking at day 1 (cluster 1) are enriched in enterocytes, whereas genes that peak at day 6 (cluster 6) and more prominently at day 8 (cluster 8) are enriched in ISCs (Fig 3h), suggesting that our analysis traces all the way back through intestinal development. Moreover, DCM-TM detects significantly more genes than detected by scRNA-seq, which misses lowly expressed genes due to limited sensitivity (Islam et al., 2014). Using immuno-cytochemistry, we confirmed exclusive expression of cluster 1 specific proteins (SGLT1, SLC43A2) in the villus. Furthermore, cluster 6 and 8 proteins (GNL3 and NUP54) were expressed in the crypt and could be verified as novel gene markers for ISCs (Fig 3i-j, Sup Fig 4g). Genes with a maximum temporal signal at day 2 displayed the highest expression level and were expressed in most single cells across different cell types, indicating that this cluster mostly represents ubiquitously expressed genes (Fig 3h, Sup Fig 4h). These results further confirm that the DCM signal is propagated in S-phase and that DCM-TM can be used to retrieve gene activity maps retrospectively.

### Enhancer activity dynamics in ISC differentiation

We next explored whether enhancer activity could be tracked across ISC to enterocyte differentiation using ChIP- and ATAC-Seq data generated in epithelium isolated from the villus (Saxena et al., 2017). We found a clear correlation between H3K27ac, which marks active promoters and enhancers (Calo and Wysocka, 2013), DNA accessibility and DCM methylation (Sup Fig 5a,d)(Saxena et al., 2017). 42% of these H3K27ac peaks were enriched for DCM (with 80% of the high-DCM cluster labelled), whereas the majority of the remaining enhancer peaks lacked sufficient DCM sites for high confidence analysis of their state (Fig 4a, Sup Fig 5b-d). Interestingly, DCM-TM detected a limited number of bivalent enhancers marked by both H3K27ac and H3K9me3 suggesting Rpol2B is in rare cases recruited to poised enhancer states. Nevertheless, most intergenic DCM labelled DMRs are marked by active enhancer chromatin likely reflecting active intestinal enhancers (Sup Fig 5a). DCM methylation was evident at known enhancers near enterocyte (*Fabp1, Cbr*), ubiquitous (*Actb*), and ISC (*Olfm4, Znhit3*) genes peaking at different time points after the start of doxycyclin treatment (Fig 4b, Sup Fig 6b, Sup Table 3) (Kaaij et al., 2013). We annotated 51,779 intergenic DCM DMRs (> 1kb from TSS) between - dox and +dox (all stages). Clustering of these intergenic DMRs based on their temporal peak values highlighted the dynamic behaviour of enhancer activity during cell state specification (Fig 4c). As expected, enrichment of H3K27ac at DMRs, was more pronounced at early than later time points as these DMRs reflect enterocyte specific and ubiquitous enhancers that are active in villi (Fig 4c). Interestingly, comparison of the different enhancer clusters based on peak day with ATAC-seq data indicates that enterocyte enhancers are accessible, whereas enhancers active at earlier stages of differentiation lose accessibility in enterocytes. We did not observe this dynamic behaviour in accessibility for TSSs, (Sup Fig 6b). Density analysis of enhancers of the different clusters around the different gene clusters showed a coordination in peak days of enhancers and nearby genes in the enterocyte differentiation process (Fig 4e, Sup Fig 6c). Genes displaying a maximum at day 2 display different enhancer kinetics as this cluster consists of both ubiquitously expressed and stage specific genes (Sup Fig 6d). ChromVAR motif analysis on enhancer regions confirmed enrichment of motifs for ISC specific transcription factors (TFs) peaking at day 8 (TCF4 and TEAD1). Enterocyte differentiation associated TFs, including ELF3, KLF5 as well as HNFA/G, known to have a crucial role in enterocyte differentiation, peak at early time points (Fig 4f, Sup Fig 6e, Sup Table 4)(Chen et al., 2019; Ng et al., 2002). In addition, by selecting TFs displaying coordinated timing of gene body and enhancer DCM labelling (Pearson r>0.3) we were able to identify *Mef2b* and *Tgif1* as potentially novel candidate TFs in ISC homeostasis and enterocyte differentiation similar to their proposed roles in other systems (Fig 4g)(Ito et al., 2017; Lee et al., 2015). In addition, reverse-coordinated peak timing (Pearson <-0.3) with maximum DCM gene body labelling in ISCs and maximum motif labelling in enterocytes identified known and putative new repressors (*Atf7*, *Glis2*, and *Mixl1)* of the ISC state (Liu et al., 2019). These results show that DCM-TM can be utilized to detect enhancer activity and relate these to underlying transcription factor dynamics.

**Figure 5.**
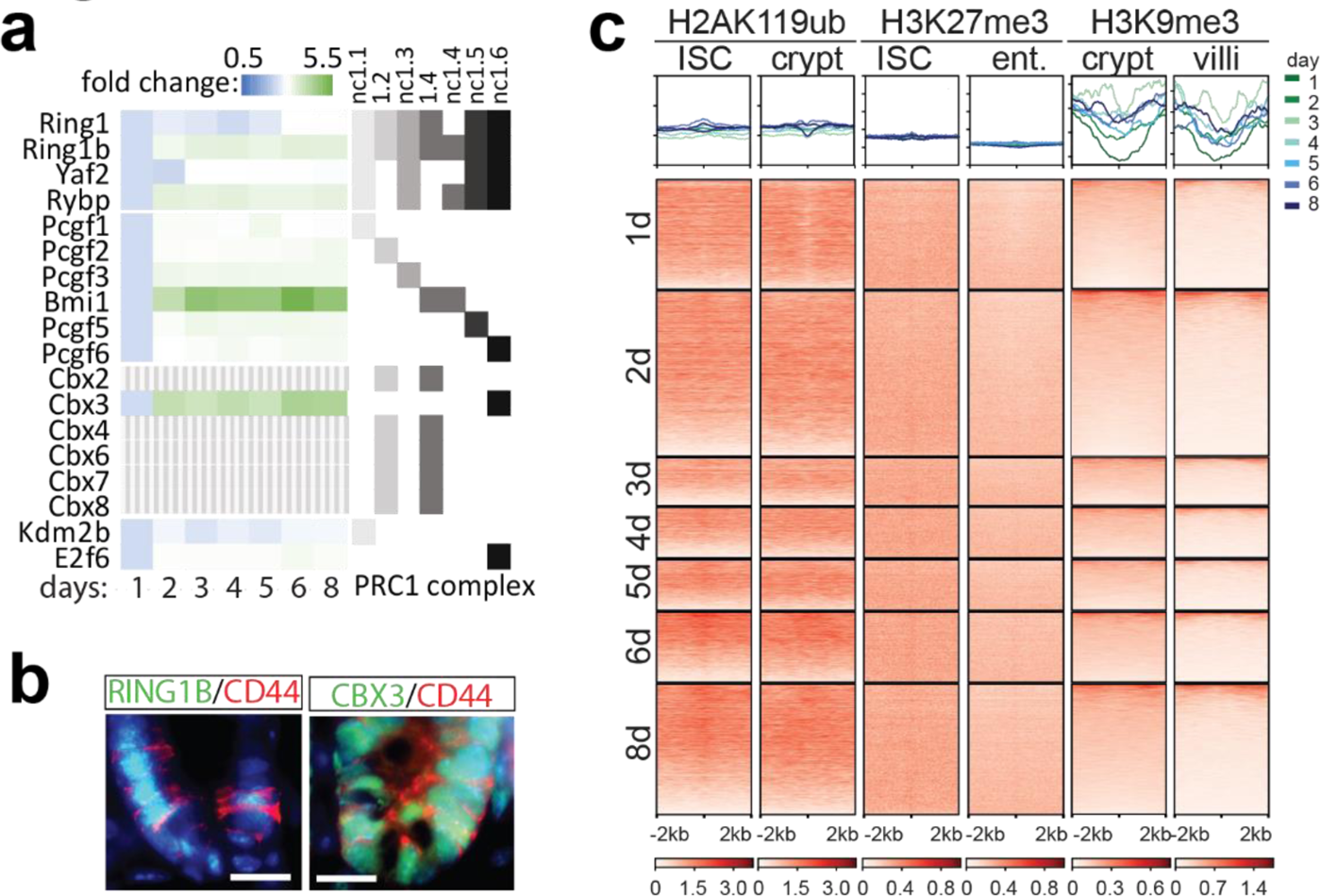
Lack of heterochromatin dynamics at intestinal enhancers. (a) Temporal behaviour of DCM methylation (normalized to t=1 day) of members of different PRC1 complexes (genes indicated in dashed grey do not accumulate DCM signal above background). (b) Immunofluorescence detection of RING1B (FITC), CBX3 (FITC) and CD44 (Texas Red) in the intestinal crypt (DNA=DAPI, scale bar: 16μm). (c) Heatmap showing H2A119ub, H3K27me3 and H3K9me3 ChIP-seq overlap in indicated cell types with the regions around enhancer DMRs peaking at different days of dox induction. Each profile plot has the same y-axis range as its corresponding heatmap.

**Figure 6.**
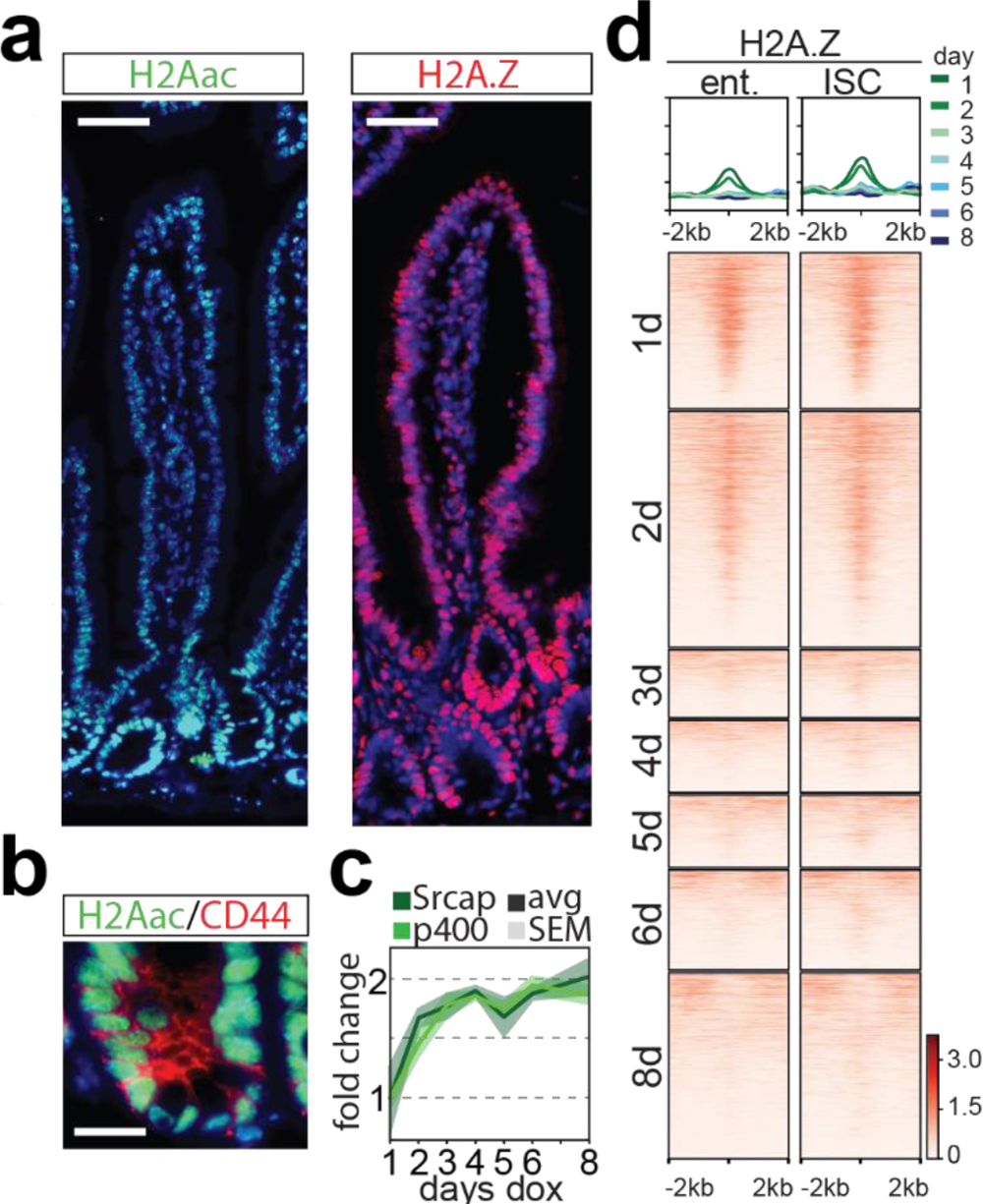
H2A.Z is recruited to enterocyte specific enhancers in ISCs. (a) Immunocytochemistry detecting H2Aac (FITC) and H2A.Z (Texas Red) (DNA=DAPI, scale bar: 50μm), and (b) combined detection of H2Aac (FITC) and CD44 (Texas Red) in the intestinal crypt (DNA=DAPI, scale bar: 16μm). (c) DCM labelling (fold change in DCM reads relative to total and normalized to T=1d) of *Srcap* and *P400*. (d) Heatmap showing H2A.Z ChIP-seq overlap in enterocytes and ISC with the regions around enhancer DMRs peaking at different days of dox induction. Enhancers were ordered according to H3K27ac and ATAC-seq enrichment (fig 4d). Each profile plot has the same y-axis range as its corresponding heatmap.

### ISC gene expression dynamics is independent of polycomb mediated repression

DCM-TM revealed *Bmi1,* a member of non-canonical polycomb complex PRC1.4, to be expressed in the ISC (Figure 3c and f). The family of PRC1 complexes is formed around a core of RING1A/B that interacts with PCGF partner proteins to form canonical PRC1 (PCGF2) and non-canonical PRC1 complexes (PCGF1, PCGF3, BMI, PCGF5, PCGF6, Fig 5a) involved in the formation of facultative heterochromatin through deposition of H2A119ub (Aranda et al., 2015). In the intestine studies involving loss of PRC1 members *Bmi1* or *Ring1b* suggested that PRC1 is required for ISC maintenance by preventing the ectopic expression of non-lineage genes through active repression (Chiacchiera et al., 2016a; Lopez-Arribillaga et al., 2015). Indeed DCM-TM indicated several other members of PRC1 complexes including *Ring1b* and *Cbx3* to show maximum gene body DCM labelling on days 6 and 8, and enrichment of RING1B and CBX3 was confirmed in CD44 positive ISCs (Fig 5b, Sup Fig 7a). In contrast, DCM gene body labelling of canonical PRC1 members *Cbx2,4,6,7* and 8 was below background levels, indicating that involvement of PRC1 in ISC maintenance is mediated through ncPRC1 complexes (Fig 5a)(Blackledge et al., 2014). To investigate the role of PRC1 and PRC2 in promoter and enhancer regulation during ISC to enterocyte differentiation, we examined enrichment of H2A119ub or H3K27me3, catalysed by PRC1 and PRC2 respectively, using ISC, crypt and enterocyte ChIP-seq data sets (Chiacchiera et al., 2016a; Ferrari et al., 2021). This analysis revealed a lack of enrichment of both H2Aub119 and H3K27me3 at enhancers at any stage of ISC differentiation (Fig 5c). Similarly, these modifications were excluded from TSSs at all stages of ISC differentiation (Sup Fig 7b).

**Figure 7.**
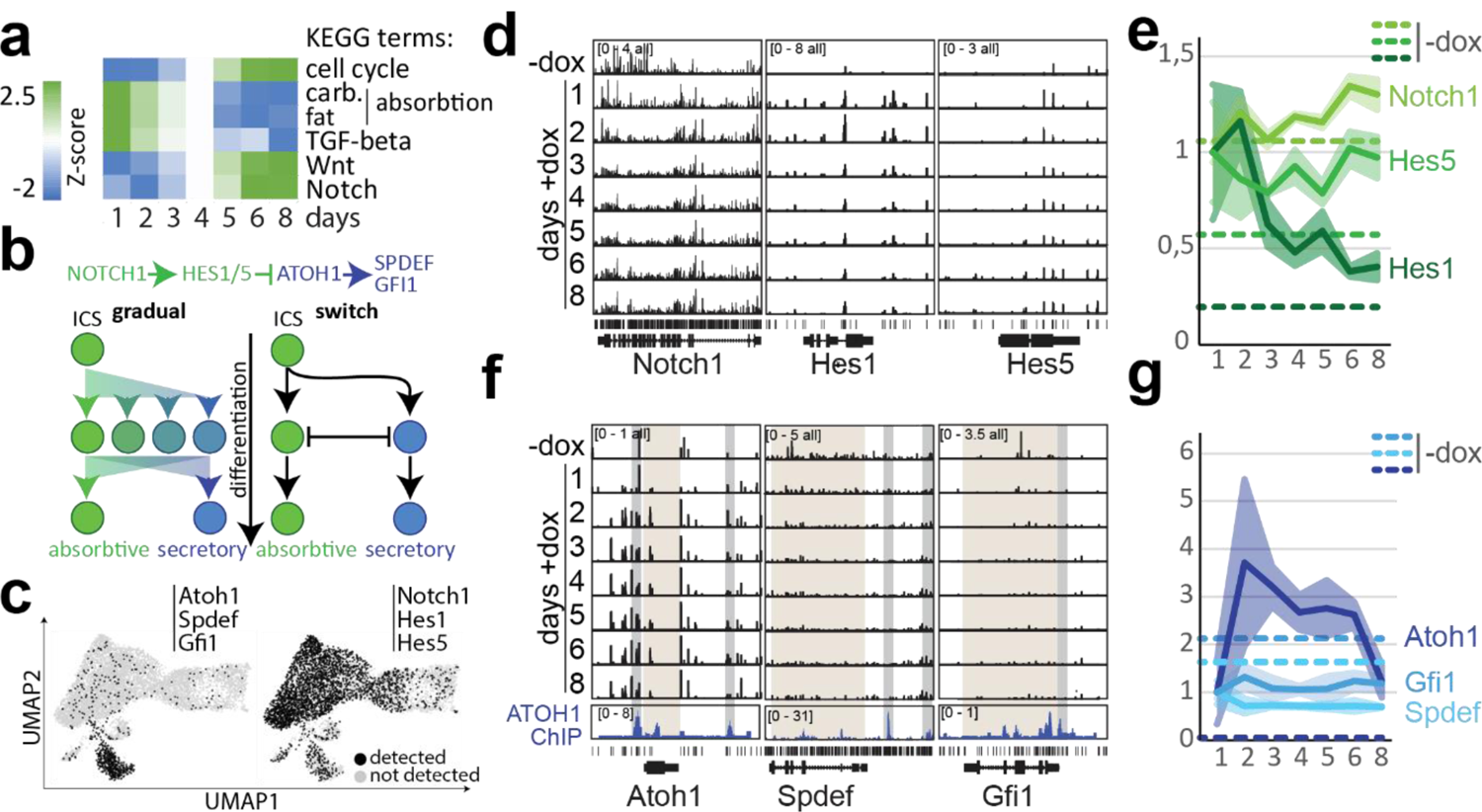
Notch signalling in absorptive versus secretory cell fate decision. (a) KEGG pathway enrichment analysis at different time points after start of dox treatment. (b) Notch signalling pathway in intestinal stem cell differentiation towards absorptive and secretory lineage; two possible mechanisms involving a gradual change or an on/off switch in Notch signalling are shown. (c) Cells expressing at least one of following three genes; *Atoh1*, *Spdef*, and *Gfi1* (left) or *Notch1*, *Hes1* and *Hes5* (right) in scRNA-seq shown in UMAP of Figure 4g. (d,e) Genome browser view of MeD-seq DCM reads in *Notch1* and its target genes *Hes1* and *Hes5* (d), and quantification of DCM signal normalized to day 1 (dashed lines represent -dox signal per gene). (f,g) As in (e,f) but now for *Atoh1*, *Spdef* and *Gfi1.* Bottom tracks in (f) show ATOH1 ChIP-seq signal from *Atoh1* GFP+ cells from the small intestine (ATOH1 targeted regulatory regions are indicated in grey and gene bodies in brown).

Several *Hp1* associated factors involved in maintenance of constitutive heterochromatin also belonged to clusters 6 and 8 (Sup Fig 7c). Nevertheless, only moderate enrichment of its target H3K9me3 (and no enrichment of H3K9me2) was detected in the crypt (Sup Fig 7d). Similar to PRC1/2 mediated histone modifications, analysis of published H3K9me3 ChIP-seq data did not reveal enrichment of H3K9me3 on enhancers and genes active at different stages of the ISC to enterocyte differentiation (Fig 5c, Sup Fig 7b). These results suggest that chromatin modifying complexes PRC1, PRC2 and HP1 complex members are enriched in the ISC but challenge the notion that they regulate differentiation associated genes through active repression of promoter and enhancer sequences.

### H2A.Z is loaded on enterocyte specific enhancers in ICS

The absence of heterochromatin mediated regulation of intestinal specific enhancers and promoters suggests that activation signals and TF networks may play a dominant role during ISC differentiation. Histone variant H2A.Z has been implicated in lineage specific gene activation (Giaimo et al., 2019). H2A.Z incorporation is mediated by *Srcap* and *P400*, which both peak on days 6 and 8, and is preceded by H2A acetylation, which showed marked enrichment in ISCs whereas HA2.Z is more uniformly distributed in the crypts and villi (Fig 6a-c). Using ISC and enterocyte specific ChIP-seq data we found that H2A.Z accumulation at ISC and enterocyte specific TSSs (Sup Fig 8a)(Kazakevych et al., 2017). Interestingly, enhancers that become active in enterocytes peaking on day 1 or day 2 showed enrichment of H2A.Z both in ISCs and enterocytes (Fig 6d). No H2A.Z enrichment was observed at ISC specific enhancers peaking on day 6 or day 8, indicating that H2A.Z pre-marks enhancers in ISCs that are poised for activation in enterocytes, and suggest a distinct role for H2A.Z in enhancer and gene activity regulation. HOMER motif enrichment analysis of H2A.Z enhancer peaks present in enterocytes revealed enrichment of TF binding sites for factors involved in Notch signalling (RBPJ and SPDEF) and several targets of the EGF signal transduction pathway including ELK1, ELK4, cMYC, and cJun, suggesting a role for these pathways in H2A.Z recruitment (Sup Fig 8b). These findings emphasize the presence of an ISC-specific chromatin landscape to maintain ISC stemness and lineage identity, and to prepare and delineate enterocyte specific enhancers and genes for future activation upon cell differentiation.

### The absorptive-secretory switch in ISC differentiation

To better understand cell state changes in enterocyte differentiation we performed KEGG pathway analysis on DCM-labelled genes and found enrichment of pathways including absorption and TGF-beta signalling at early time points. Pathways including cell cycle, Wnt, EGF and Notch signalling showed enrichment at later time points, consistent with their lineage history (Fig 7a, Sup Table 5). Notch signalling controls the absorptive versus secretory cell fate decision dictating repression of *Atoh1* in the ISC and enterocyte progenitors through action of *Hes1*,*3* and *5* (Fig 7b). Loss of contact of proliferating ICSs (Notch+) with Paneth cells expressing the Notch ligand *Dll1* leads to downregulation of *Notch1* and subsequent upregulation of *Atoh1* in future secretory cells (Beumer and Clevers, 2020).

Notch mediated repression of *Atoh1* and its key target genes *Spdef* and *Gfi1* could involve a gradual transition from a bi-potential progenitor to one cell state or may involve a binary switch towards the absorptive or secretory lineage. Single cell RNA-seq data demonstrated predominant expression of *Atoh1*, *Spdef* and *Gfi1* in secretory cell types, but also revealed several cells that appear committed to the absorptive lineage to express at least one of these genes (Fig 7c). In addition, a few cells express both *Notch* and *Atoh1* or don’t express *Notch* and *Atoh1* at all, making it difficult to discern how the absorptive-secretory switch is mediated (Sup Fig 8c). Examination of DCM-TM data indicated that *Notch1* as well as Notch target genes including *Hes1, Hes3* and *Hes5* are expressed throughout enterocyte differentiation (Fig 7d,e). In contrast, DCM methylation in gene bodies of *Spdef* and *Gfi1* never got above background levels (Mann-Whitney significance test p > 0.05), suggesting their activation results in irreversible commitment towards the secretory lineage. These results demonstrate that, unlike the transition from the ISC into the absorptive state, the switch towards a secretory state represents a binary one directional switch rather than a gradual transition of, transcriptional programs (Fig 7f,g).

## Discussion

To facilitate whole genome cell state tracing, we developed a system to epigenetically tag transcribed genes to be examined at later stages of development or differentiation in vivo. We applied this DCM-TM technology to perform whole transcriptome and enhancer activity lineage tracing of specific cell types, and demonstrated the possibility to establish TFs and signal transduction roadmaps through isolation of a differentiated cell type without the need to isolate progenitor or stem cells and without the need to infer connectivity in silico. We identified novel marker genes for different cell states and provide new insights into the transcriptional dynamics during cellular differentiation in the mouse intestine.

Key parameters for an epigenetic lineage tracing system to work are the application of an epigenetic tag that is normally absent in mammalian cells, is maintained and propagated upon DNA synthesis, and does not interfere with gene expression. Our study showed that DCM methylation approaches these criteria simultaneously. DCM methylation is only present at low levels in wildtype ES cells and intestinal epithelium (2-3%), while only a 5-fold and 25-fold induction in ES cells and intestinal epithelium, respectively, is sufficient to reliably identify active genes and enhancers and trace their activity back in time. In contrast to other forms of bacterial methylation which are not propagated such as DAM, propagation of DCM methylation is 78% in the intestinal epithelium (van Steensel and Henikoff, 2000). This is lower than previously described, but sufficient to detect gene body and enhancer labelling over at least 10 cell divisions (Clark et al., 1995). Lastly, we found that only a limited number of genes is affected by induction of the fusion protein. This lack of interference with transcription might be related to the fact that gene bodies of active genes already accumulate CpG methylation, thought to repel intragenic initiation of RNA pol2 (Neri et al., 2017). In addition, as the DCM motif is found much less frequently in genes it may thus have only limited effect on transcription.

In this study we applied DCM-TM to understand the mechanism directing the absorptive-secretory switch in the intestinal epithelium involving the Notch signalling pathway. Activation of *Notch* and its downstream *Hes* family targets is mediated by cell-cell contact through direct contact of Notch and its ligand Dll. Notch signalling is required for maintenance of the ISC where Notch ligands are expressed by Paneth cells, but also during ISC differentiation to consolidate the absorptive lineage (VanDussen et al., 2012). In the secretory lineage Notch is downregulated resulting in de-repression of *Atoh1* and its downstream targets *Gfi1* and *Spdef*. Our DCM-TM data demonstrates that Notch signalling remains active throughout enterocyte differentiation and that *Spdef* and *Gfi1* are never activated. This indicates that the absorptive-secretory lineage fate decision involves a committed rather than a temporarily dynamic absorptive-secretory intermediate state. Nevertheless, *Atoh1* is active in ISCs and cells that commit to the absorptive lineage, probably as a consequence of fluctuating Notch activity and to be able to quickly respond when *Notch* levels decrease below a specific threshold for locking in the secretory state. This finding explains why *Atoh1* lineage tracing studies revealed *Atoh1* positive cells to contribute to the ISC pool (Ishibashi et al., 2018), and also indicates that this *Atoh1* expression level is too low to activate *Spdef* and *Gfi1* and the downstream secretory program.

Our study and studies of others indicate that the role for complexes in heterochromatin formation and maintenance in ISC differentiation associated gene expression dynamics is very limited. Previous studies involving loss of PRC1 members *Bmi1* or *Ring1b* suggested that PRC1 is required for ISC maintenance by preventing ectopic expression of non-lineage genes (Chiacchiera et al., 2016a; Lopez-Arribillaga et al., 2015). Application of DCM-TM revealed that PRC1 mediated gene repression is mediated by non-canonical PRC1 complexes that contain RYBP which catalyses H2A ubiquitination independent of PRC2. This is in line with the observation that loss of PRC2 and H3K27me3 does not affect H2A119ub in the intestine (Chiacchiera et al., 2016b). The present study also revealed a static landscape of repressive chromatin modifications H2A119ub, H3K27me3 and H3K9me3 in intestinal homeostasis. Similarly, DNA methylation changes were found to be very limited between different epithelial cell types (Kaaij et al., 2013), suggesting that the main role of facultative and constitutive heterochromatin in ISCs is dictating repression of non-lineage genes. Therefore, activation and repression of enhancers and genes in intestinal homeostasis appears to be regulated by other epigenetic mechanisms and transcription factor networks. We found histone variant H2A.Z to be loaded on enhancers in ISCs that are destined to become activated in enterocytes, indicating that the ISC dictates and limits enhancer activity in its decedents through H2A.Z recruitment to enhancers. The clear enrichment of motifs of EGF and Notch regulated transcription factors in these enhancers make both signalling pathways the likely candidate signal for H2A.Z recruitment.

Several cell state tracing technologies detecting the history of gene expression have been described before. These include CRISPR spacer mediated recording of DNA or RNA to monitor complex cellular behaviour retrospectively, as well as smFISH based detection of Crispr/Cas mediated targeted disruption of expressed recording elements (Frieda et al., 2017; Schmidt et al., 2018). Unfortunately, all these technologies are restricted by a limited number of genes that can be recorded. The recent developments in scRNA-seq provides alternative means to detect cell states and gene expression changes in relation to developmental trajectories. Temporal changes in abundance of spliced and un-spliced gene products (La Manno et al., 2018), and minimum spanning tree analysis (Monocle) have been applied to predict cell trajectories in silico (Trapnell et al., 2014), but these analyses are limited by the temporal resolution and the number of genes detected, and therefore often fail to detect the changes in gene expression from one cell state to the next, and are more difficult to apply along developmental trajectories. The present DCM-TM technology circumvents these issues providing a genome wide picture of gene and enhancer activity at any timepoint during development or differentiation. The DCM-TM transgene can be combined with conventional lineage tracing technologies to fine map cell fate decisions, discriminate between lineage paths and keep track of network changes. In addition, DCM-TM can be applied to follow embryonic development and tissue regeneration, providing a powerful system to identify temporal maps of transcription factor networks and signal transduction pathways that can be used to improve stem cell expansion and cell differentiation models.

## Acknowledgements

We would like to thank Guillaume Jacquemin and Silvia Fre for providing protocols for enterocyte isolation, Jonathan Klaver for help, Riccardo Fodde and Joana Carvalho Moreira de Mello and department members for helpful discussions.

## Author contributions

R.G.B., J.B.B, B.T, M.C. and J.G. conceived and performed the experiments and data analysis. E.W., E.S, W.IJ., E.M. and Y.H. assisted in immunocytochemistry, FACS analysis, sequence analysis and modelling. J.L. E.G.S. and W.B. aided in development of DCM-TM and interpreting the results, and M.L. helped with motif analysis. All authors discussed the results and contributed to the final manuscript.

---

**Supplementary Figure 1.**
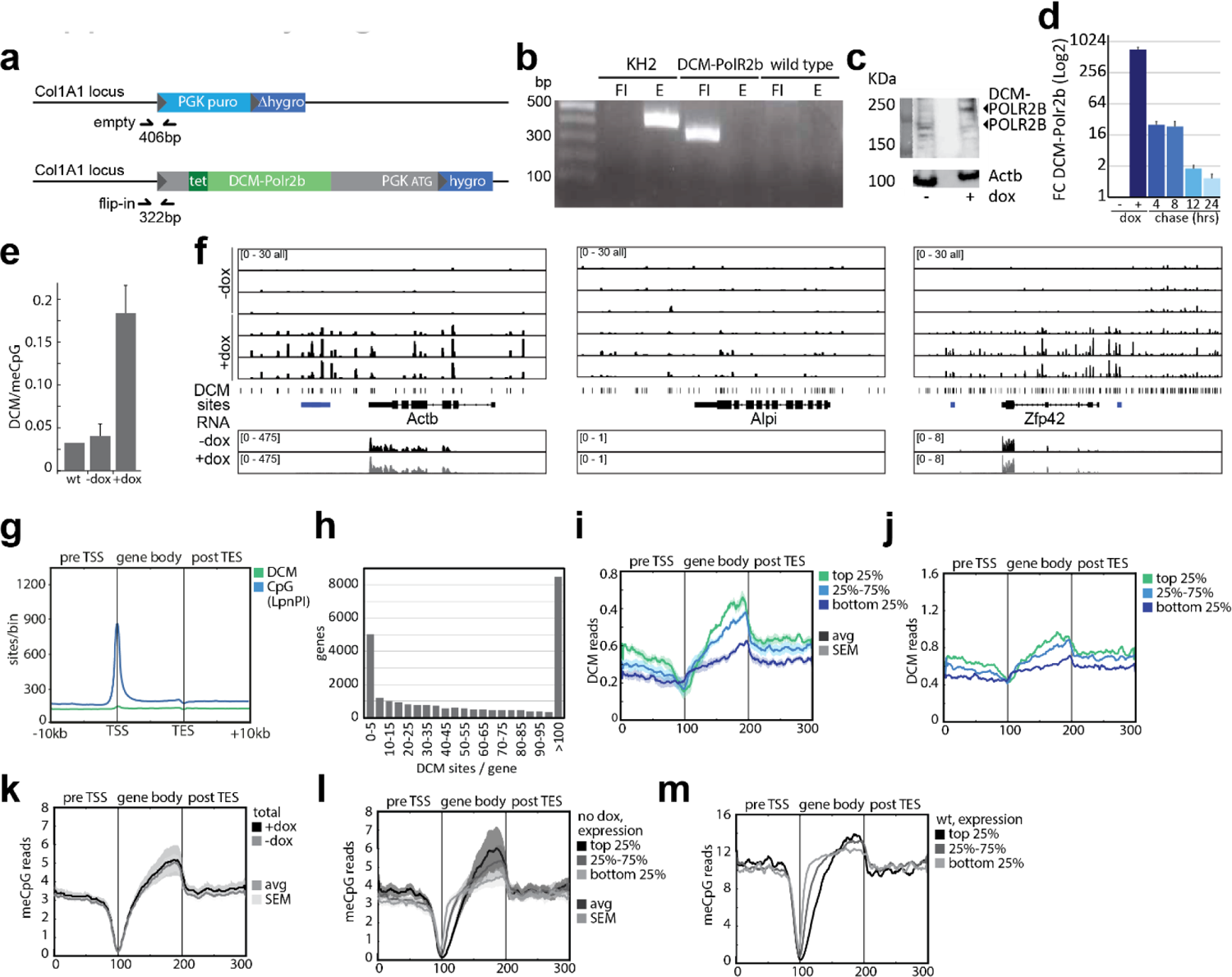
(a) The DCM-Polr2b fusion gene was introduced in the Col1A1 locus by Flipase mediated insertion. (b) PCR with primers for flip-in (FI) and empty (E) indicated in (a) verifying proper integration of the transgene. (c) Western blotting analysis detecting POLR2B, DCM-POLR2B and ACTB in uninduced and induced ES cells. (d) qRT-PCR analysis detecting DCM-POLR2B transcript in ES cells at different timepoints (hours) after removal of dox. (e) The ratio of DCM/CpG methylation genome wide in wild type (WT), DCM-Polr2b -dox and +dox ESCs. (f) Genome browser view of DCM specific MeD-seq reads in the *Actb*, *Alpi* and *Zfp42* loci (intergenic DMRs are indicated in blue). (g) Gene meta-analysis showing binned distribution of DCM and CpG sites. (h) Histogram showing distribution of DCM sites per gene. (i) Gene meta-analysis showing binned distribution of DCM reads in the top 25%, 25%-75% and bottom 25% expressed genes in DCM-Polr2b ESCs in the absence of dox. (j) As in (i) but now for wild type ESCs (n=1). (k) Gene meta-analysis showing binned distribution of all CpG reads before and after dox treatment (5 days). (l) Gene meta-analysis showing binned distribution of CpG reads in the top 25%, 25%-75% and bottom 25% expressed genes in DCM-Polr2b ESCs in the absence of dox. (m) As in (l) but now for wild type ESCs (n=1).

**Supplementary Figure 2.**
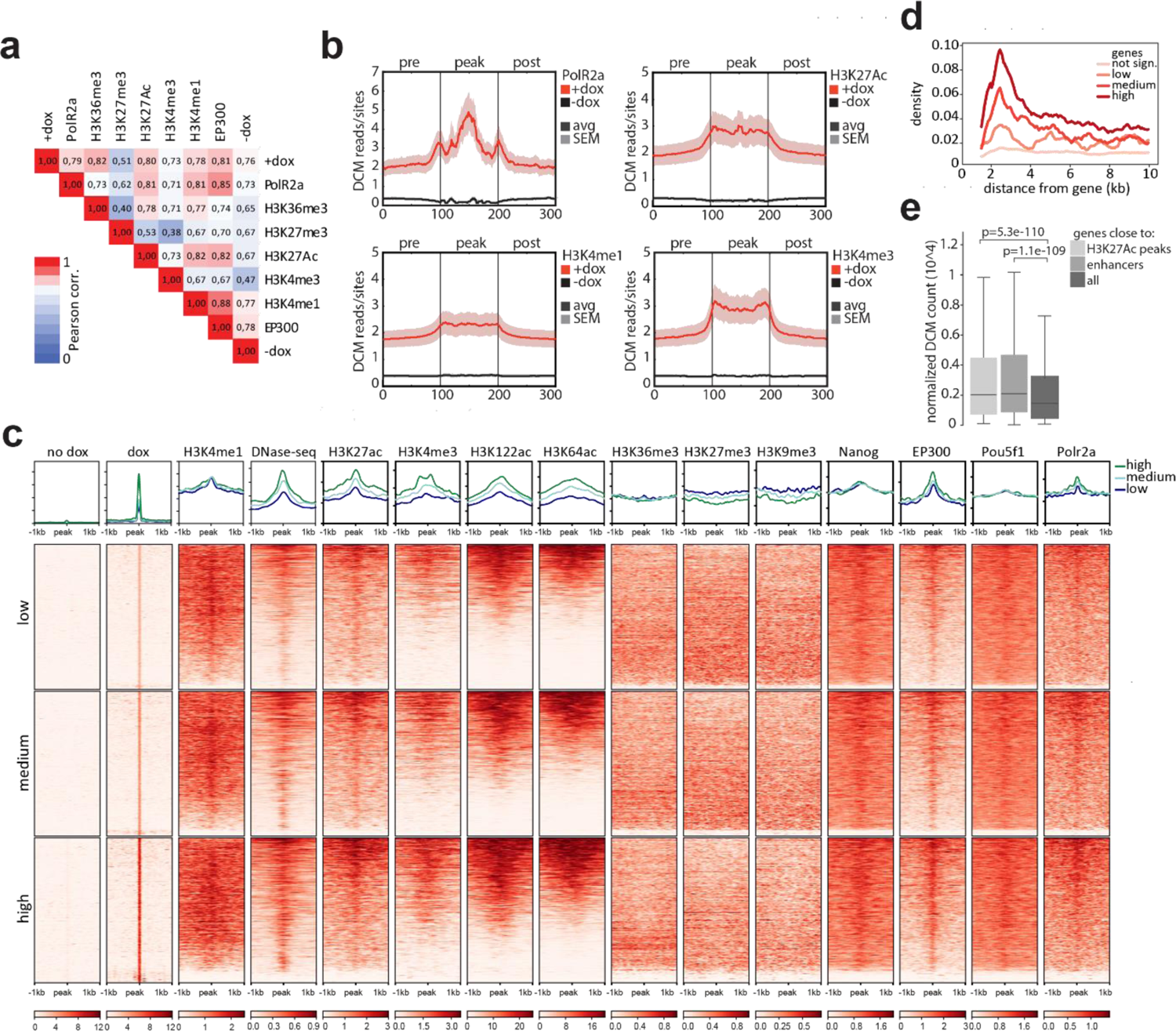
(a) Pearson correlation analysis comparing DCM and ChIP-seq read count distribution. (b) Enhancer meta-analysis showing binned distribution of DCM read counts over Polr2a, H3K27Ac, H3K4me1 and H3K4me3 positive genomic regions and 1kb proximal and distal flanking regions. (c) Heatmap showing ChIP-seq overlap with the regions around enhancer DMRs. DMRs are split in three equal clusters based on the normalized number of reads for +dox. Each profile plot has the same y-axis range as its corresponding heatmap. (d) Density plot showing the number of enhancer DMRs in the 10kb region around genes that were either not significantly labelled by DCM or genes split in three equal clusters based on fold change between +dox and –dox. (e) Normalized DCM count of +dox samples for genes close to the enhancer DMRs, close to H3K27ac peaks and all genes. P-values were calculated using a one-sided Wilcoxon rank-sum test.

**Supplementary Figure 3.**
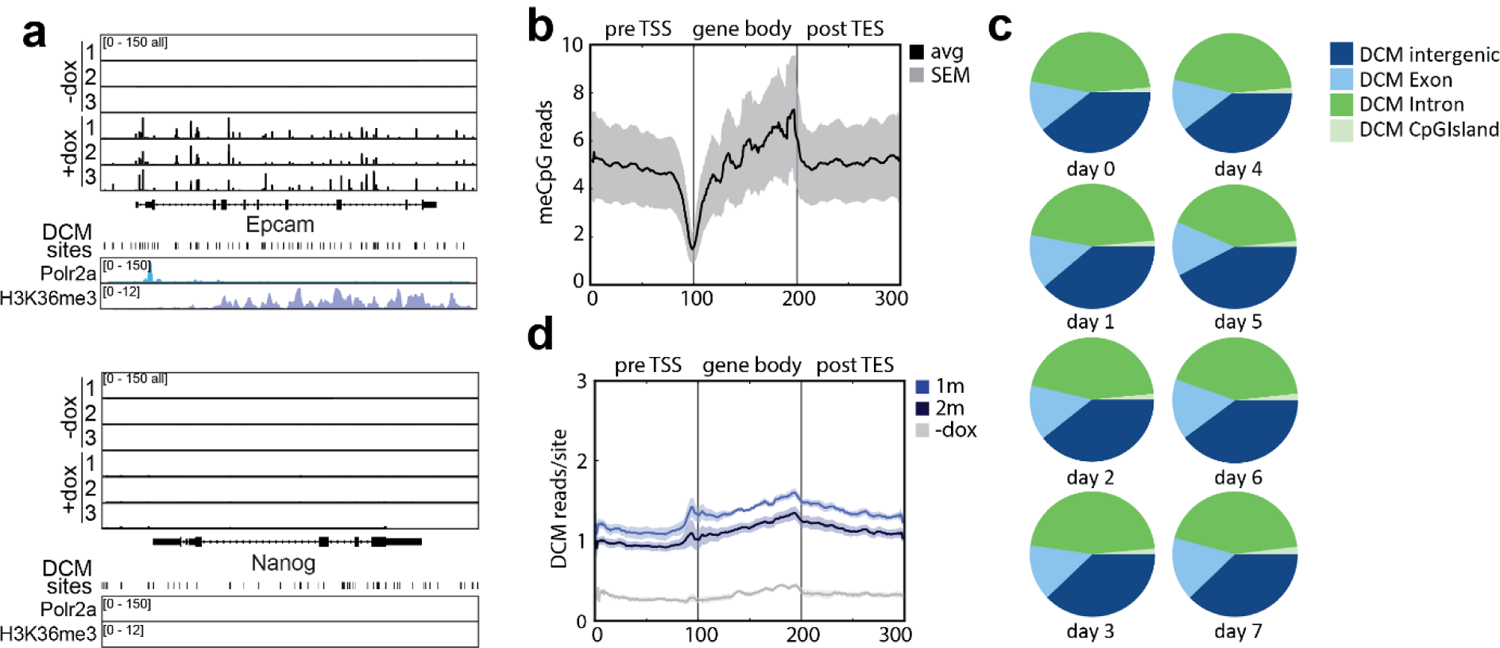
(a) Genome browser view of *Epcam* and *Nanog* loci showing DCM specific MeD-seq reads before and after 1 day dox treatment. POLR2a and H3K36me3 ChIP-seq tracks are shown below. (b) Gene meta-analysis showing binned distribution of all CpG reads in the absence of dox. (c) Relative distribution of DCM reads in intergenic, exonic, intronic and CpG island sequences at indicated days after removal of doxycycline. (d) Gene meta-analysis showing binned distribution of DCM reads in -dox control mice and mice treated for 7 days with dox followed by a 1 and 2 month chase prior to tissue harvesting.

**Supplementary Figure 4.**
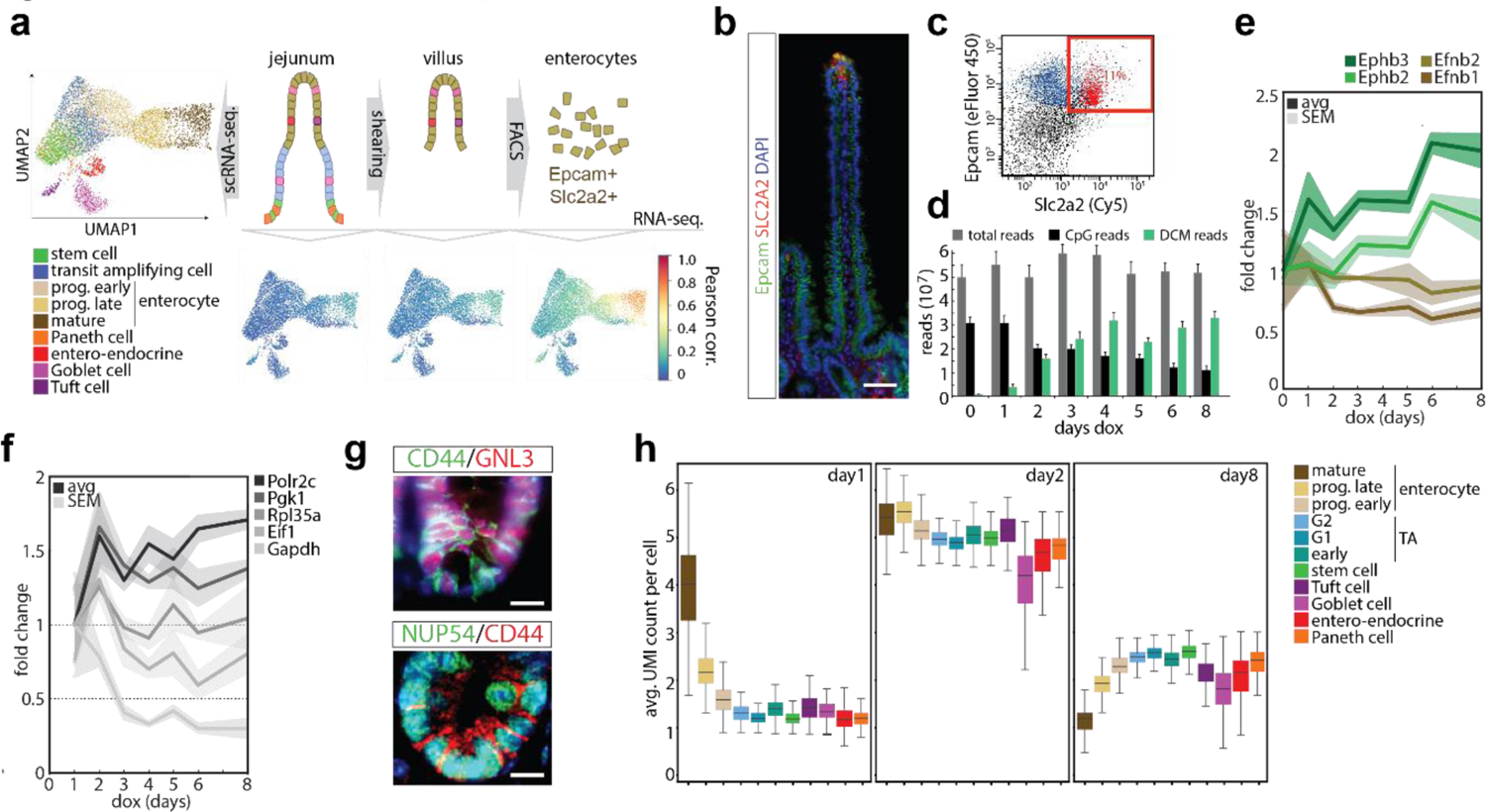
(a) Overview of the experimental procedure to isolate SLC2A2 expressing enterocytes. Villi were isolated from intestinal epithelium of jejunum followed by FACS isolation of SLC2A2 positive cells. Left panel shows UMAP of scRNA-seq data colored according to annotation as specific cell types, bottom panels show Pearson correlation analysis of bulk RNA-seq analysis on total epithelial, villi and SLC2A2 positive fractions with scRNA-seq data. (b) Immuno-cytochemistry detecting Epcam (FITC) and SLC2A2 (Texas red, DNA in DAPI, scale bar: 50μm). (c) FACS analysis of intestinal epithelial cells, SLC2A2 and Epcam positive cells are highlighted in red. (d) Genome wide DCM and CpG methylation level at different time points after start of dox induction. (e,f) DCM labelling (relative to total and normalized to T=1d) of *Ephb2, Ephb3, Efnb1*, and *Efnb2* genes (e), and the ubiquitously expressed *Polr2c, Pgk1, Rpl35a, Eif1* and *Gapdh* genes at different time points after start of dox treatment (f). (g) Immunocytochemistry detecting GNL3 and NUP54 in combination with CD44 (DNA is DAPI, scale bar: 16μm) (h) Normalized UMI count distribution per cell type on day 1, 2 and 8 DCM labelling peak time point, showing increased expression of genes at day 2.

**Supplementary Figure 5.**
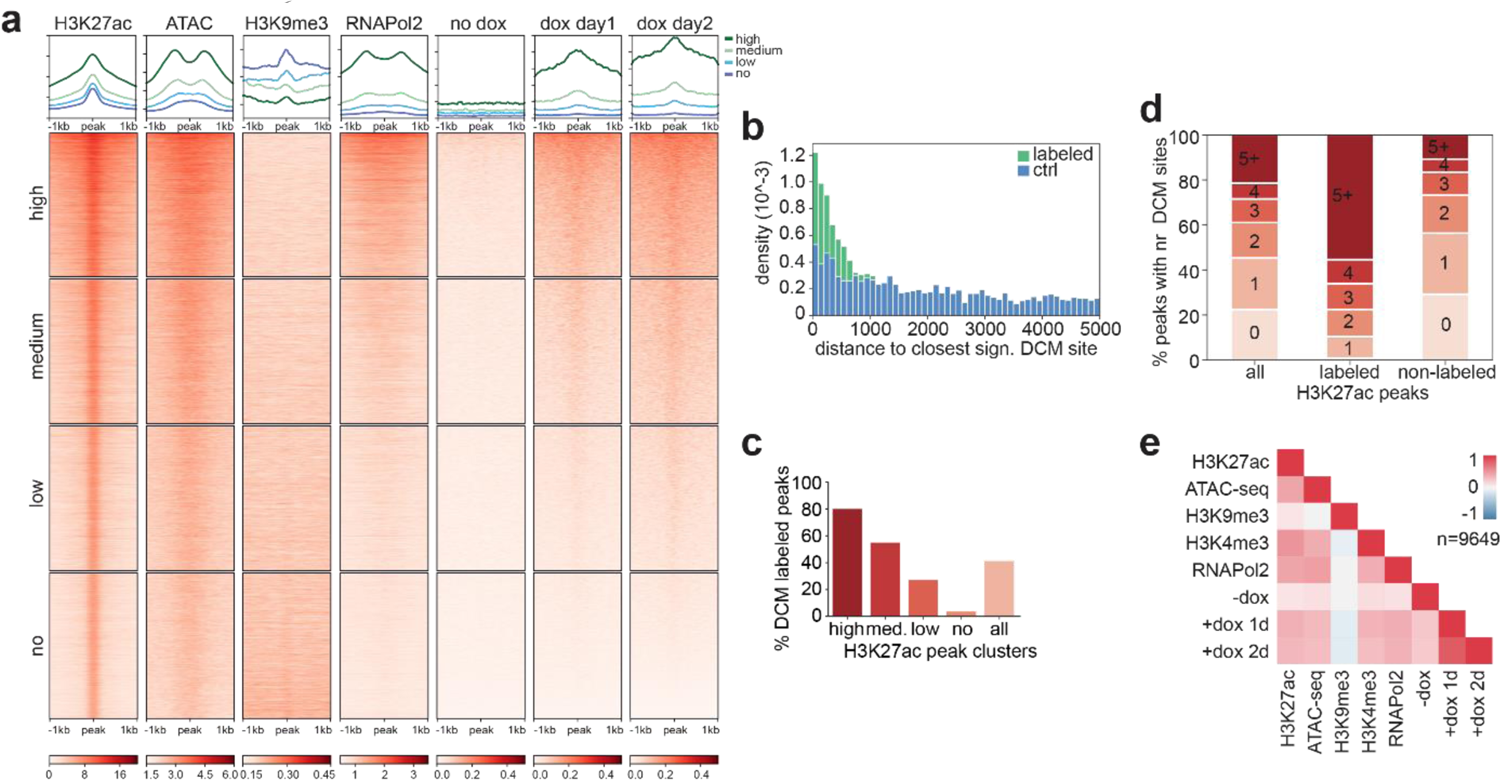
(a) Heatmap showing the overlap of villi H3K27ac peaks with several villi ChIP-seq and DCM datasets. All ChIP-seq datasets were generated from villi samples [Saxena et al. 2017] and the DCM data from – dox samples and the day 1 and day 2 +dox samples are shown. H3K27ac peaks are ordered according to overlapping DCM signal ±1kb of the peak center and split in four equally sized clusters based on this ordering. Each profile plot has the same y-axis range as its corresponding heatmap. (b) Histogram with the distance to the closest significant DCM site for both the DCM labeled H3K27ac peaks (i.e. peaks with ≥ 1 significant DCM site) and random controls based on 100 sets of reshuffled H3K27ac peaks. Density for each 100 bp bin up to 5kb is shown. (c) Percentage of H3K27ac peaks that are labelled by DCM (i.e. peaks with ≥ 1 significant DCM site <750 bp from peak). The H3K27ac peaks are split in four clusters based on H3K27ac intensity and related to all peaks. (d) Barplot showing the number of DCM sites overlapping each peak. The percentages with each number of sites are shown for all peaks, the labeled peaks (i.e. peaks with ≥ 1 significant DCM site) and non-labeled peaks (i.e. peaks wihout significant DCM site). (e) Correlation heatmap showing the spearman correlation at the H3K27ac peaks between the different ChIP-seq and DCM datasets shown in (a). The number of reads overlapping each H3K27ac peak with ≥ 3 DCM sites were normalized for the peak length or the number of DCM sites for the ChIP-seq and DCM datasets, respectively.

**Supplementary Figure 6.**
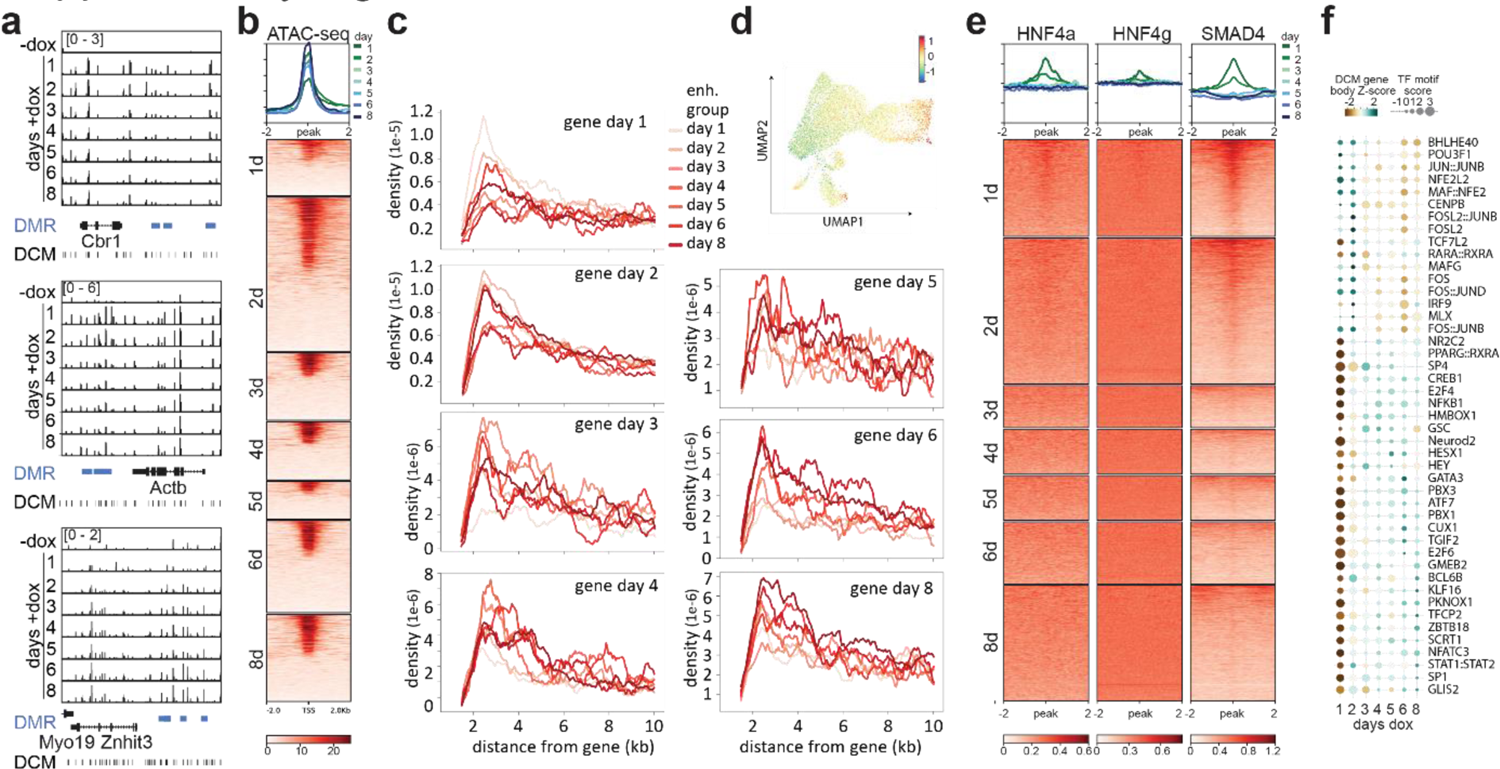
(a) Genome browser view showing DCM labelling of enterocyte specific (*Cbr1*), ubiquitous (*Actb*) and ISC specific (*Znhit3*) genes with enhancers (in blue) nearby showing similar behaviour in time. (b) Heatmap showing ATAC-seq overlap with the regions around TSS of genes peaking at different days of dox induction. (c) Density plot showing the number of enhancer DMRs per peak day in the 10kb region around genes split in clusters based on peak timing of gene body DCM labelling. (d) Differential enrichment of genes peaking on day 2 split by having relatively more enhancers peaking on day 2 or day 8 nearby; difference in gene expression plot on UMAP showing increased expression of genes linked to day 2 enhancers in enterocytes. (e) Heatmap showing ChIP-seq overlap for HNFa, HNFg and SMAD4 with regions around enhancer DMRs. Enhancers were split in clusters based on the maximum day of DCM accumulation. (f) (f) Combined analysis of motif enrichment and DCM gene body labelling dynamics of TFs displaying a negative correlation in time (right).

**Supplementary Figure 7.**
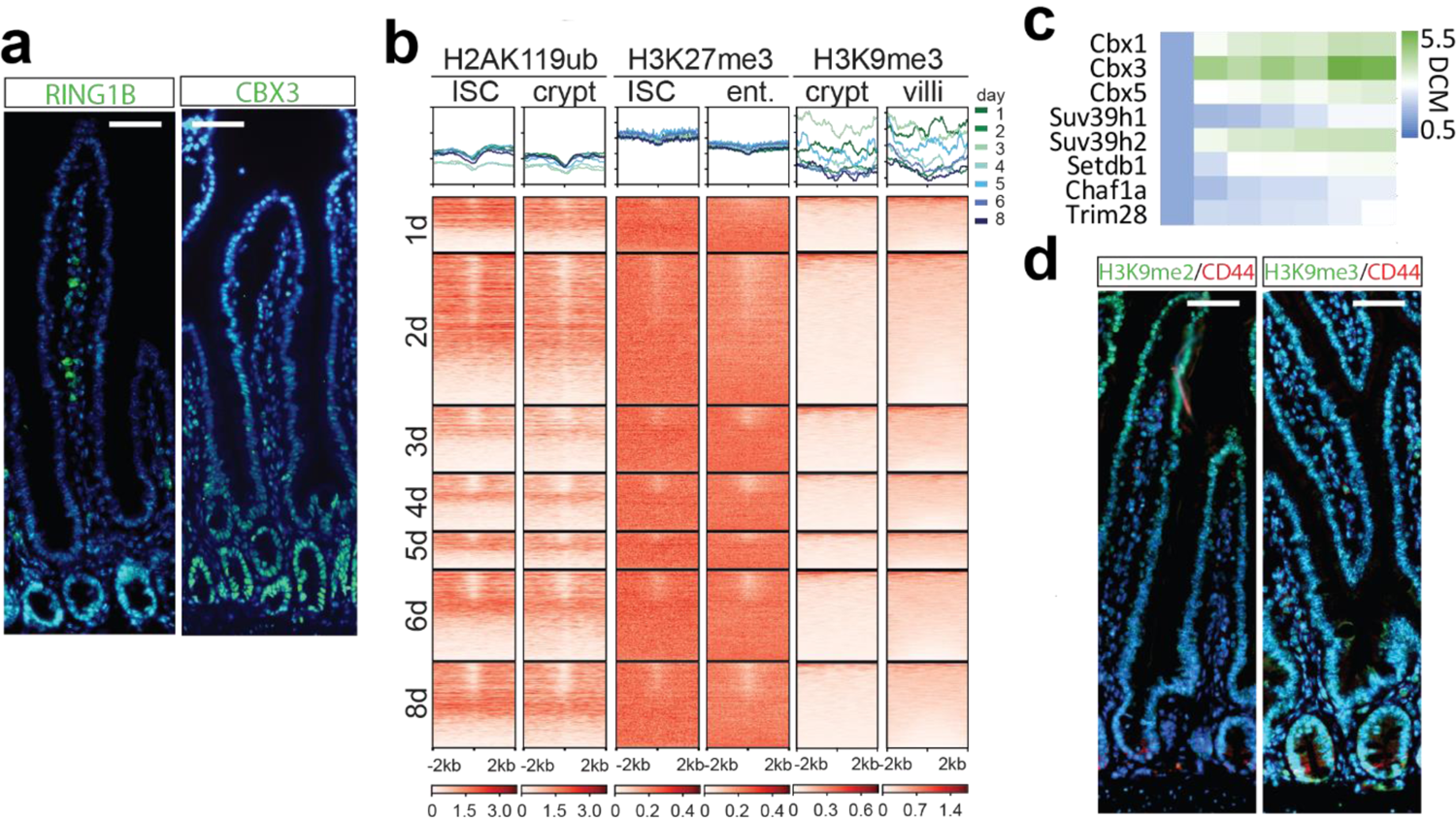
(a) Immunocytochemistry detecting RING1B and CBX3 (FITC) (DNA=DAPI, scale bar: 50μm). (b) Heatmap showing ChIP-seq overlap in enterocytes, ISC, crypt and villi for H2A119ub, H3K27me3, and H3K9me3 with regions around the TSS DMRs. TSS were split in clusters based on the maximum day of DCM accumulation and ordered according to H3K27ac and ATAC-seq enrichment. (c) Temporal behaviour of DCM methylation (normalized to t=1 day) of genes encoding proteins involved in establishment and maintenance of constitutive heterochromatin. (d) Immunocytochemistry detecting H3K9me2 and H3K9me3 (FITC) in combination with CD44 (Texas Red, scale bar: 50μm).

**Supplementary Figure 8.**
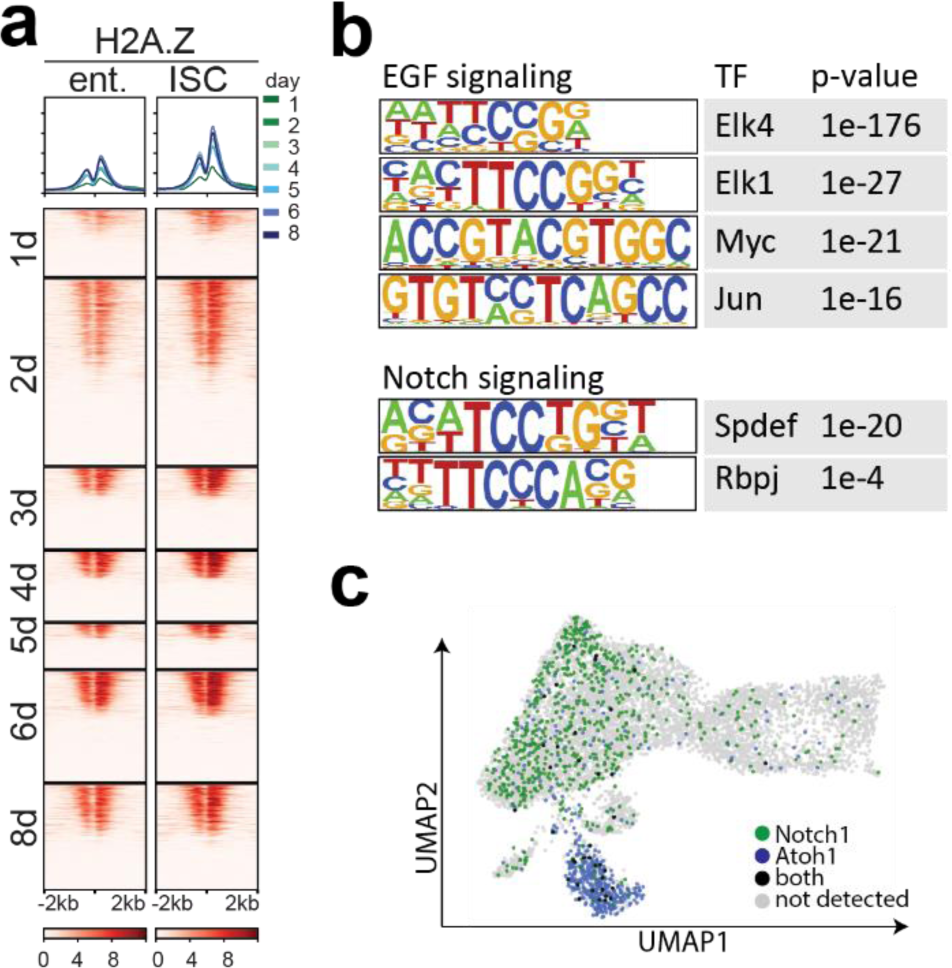
(a) Heatmap showing H2A.Z ChIP-seq overlap in enterocytes and ISC with regions around the TSS DMRs. TSSs were split in clusters based on the maximum day of DCM accumulation and ordered according to H3K27ac and ATAC-seq enrichment. (b) HOMER motif analysis on H2A.Z enhancer peaks present in enterocytes revealing motif enrichment for TFs downstream of EGF and Notch signalling. (c) UMAP displaying cells in which only *Notch1* or *Atoh1* was detected and cells where both genes or no gene was detected (plotted in UMAP shown in Figure 4g).

**Supplementary Table 1** Sequencing statistics.

**Supplementary Table 2** Overview of gene body DCM counts in uninduced and induced ES cells and enterocytes.

**Supplementary Table 3** Overview of intergenic differentially methylated DCM sites in uninduced and induced ES cells and enterocytes.

**Supplementary Table 4** Motif analysis on intergenic DMRs at different timepoints of dox induction in enterocytes.

**Supplementary Table 5** KEGG pathway analysis on differential DCM labelling of genes upon dox induction.

